# Neural Responses to Unexpected Stimulus Repetitions and Omissions in Auditory Cortex Provide Mixed Evidence for Predictive Coding

**DOI:** 10.64898/2026.03.30.715297

**Authors:** Bhanu Shukla, Harry Shirley, Liad Goodovitch, Yonatan I. Fishman, Yale E. Cohen

**Affiliations:** Department of Otorhinolaryngology, University of Pennsylvania, Philadelphia, PA 19104 USA; Departments of Neurology and Neuroscience, Albert Einstein College of Medicine, Bronx, NY 10461, USA; Departments of Otorhinolaryngology and Bioengineering, University of Pennsylvania, Philadelphia, PA 19104 USA

**Author notes:** Corresponding author: Yale E. Cohen, PhD, Department of Otorhinolaryngology, 3400 Spruce St, 5 Ravdin, Philadelphia, PA 19147.

## Abstract

Humans and other animals process sensory uncertainty by integrating stimulus information with prior knowledge and expectations. Predictive coding conceptualizes perception as a form of Bayesian inference wherein hierarchical brain circuits update internal models to reconcile bottom-up sensory input with top-down predictions. Whereas predictive coding is a leading theory, the extent to which it is implemented in primary sensory cortices remains a matter of debate. To further investigate this issue, we examined single-neuron spiking activity in macaque primary auditory cortex (A1) to expected versus unexpected stimulus repetitions and to expected versus unexpected omissions. On average, we found that A1 neurons did not show enhanced responses to unexpected stimulus repetitions, contrary to predictive-coding theory. However, they did show enhanced responses to unexpected stimulus omissions. Taken together, these mixed results place empirical restraints on how PC is implemented in A1.

**Significance Statement:** Perception depends on the brain’s ability to infer the causes of sensory inputs by integrating new information with prior knowledge under uncertainty. Our results reveal nuanced evidence for predictive coding within the primary auditory cortex (A1). Specifically, spiking activity during unexpected stimuli and unexpected stimulus omissions provide conflicting and supporting, respectively, data for this Bayesian framework. These findings refine our understanding of neural mechanisms underlying perception and provide empirical constraints on the neurobiological implementation of predictive processing.

## Introduction

Humans and other animals generate perceptual representations of the world to guide adaptive behavior despite uncertainty in the sensory input (Knill and Richards, 1996; Gold and Shadlen, 2002; Gold and Shadlen, 2007; Griffiths and Tenenbaum, 2011; Aitchison and Lengyel, 2017; Heilbron and Chait, 2018). Perception can be considered a process by which this sensory uncertainty is integrated with prior knowledge in a manner that follows the rules of Bayesian inference. In hearing, for example, such inference is crucial for learning language, parsing speech in noisy environments, and music appreciation (Denham and Winkler, 2006; Szabó et al., 2016; Zarcone et al., 2016; Koelsch et al., 2019).

One highly influential neural implementation of Bayesian inference is “predictive coding” (PC) (Rao and Ballard, 1999; Friston, 2005; Shipp, 2016; de Lange et al., 2018; Heilbron and Chait, 2018). Under the PC framework, internal perceptual models are updated based on “surprise” (i.e., “prediction-error”) responses, which are elicited when there is a mismatch between “bottom-up” sensory input and “top-down” expectations (i.e., predictions). However, there is an active debate as to whether and to what extent PC is implemented in the auditory cortex (Parras et al., 2017; Carbajal and Malmierca, 2018; Heilbron and Chait, 2018; Teichert et al., 2025). A key question in this debate is whether putative prediction-error responses can explained more parsimoniously by local bottom-up neurophysiological phenomena such as stimulus-specific adaptation (SSA), forward suppression, or “off” responses (May and Tiitinen, 2010; Fishman, 2014; Aitchison and Lengyel, 2017; Symonds et al., 2017; Heilbron and Chait, 2018; Ross and Hamm, 2020; May, 2021; O’Reilly, 2021).

Here, we expanded upon previous efforts to differentiate PC from local bottom-up mechanisms (e.g., SSA) by recording from the primary auditory cortex (A1) of macaque monkeys during auditory pattern and omission stimulus paradigms (Alain et al., 1994; Hughes et al., 2001; Wacongne et al., 2011; Wacongne et al., 2012; Fishman, 2014). During the pattern paradigm, PC predicts enhanced responses to an unexpected repetition of an auditory stimulus. In contrast, SSA predicts reduced responses. During the omission paradigm, PC predicts enhanced responses to an unexpected stimulus omission. This cannot readily be explained by SSA because SSA results from the repetition of physically presented stimuli and not the absence of a stimulus.

Contrary to PC theory, our results did not reveal reliable evidence for enhanced responses to unexpected stimulus repetitions in A1. On the other hand and consistent with PC, we found that unexpected stimulus omissions elicited stronger responses than expected ones; these stronger responses could reflect either a prediction-error signal or a neural correlate of a top-down prior expectation (Bendixen et al., 2009; Wacongne et al., 2011; Sanmiguel et al., 2013a; Kok et al., 2014; Schröger et al., 2015; Chennu et al., 2016; de Lange et al., 2018; Heilbron and Chait, 2018; Demarchi et al., 2019; Yaron et al., 2025). These mixed results place empirical restraints on how PC is implemented in A1.

## Results

We report data from 198 single units that were recorded across 60 sessions from Monkey C and 229 single units that were recorded across 53 sessions from Monkey W. In all sessions, we recorded data from at least 2 of the 3 auditory paradigms (i.e., the oddball, pattern, and/or omission paradigms). Across different recording sessions, we modified the presentation rate of the tone bursts as well as the temporal occurrence of the tone burst that violated the expectation (e.g., in some sessions, it was presented randomly within a tone-burst sequence or at regular intervals). However, because our analyses indicated that these manipulations did not systematically change our neural population’s responses to the tone burst that violated expectations, we combined the data in the presentation below.

### CSD profiles and oddball responses are characteristic of the primary auditory cortex

We identified the primary auditory cortex (A1) through a series of functional assays, including the CSD profile (Steinschneider et al., 1992; Fishman et al., 2001b; Lakatos et al., 2005; Szymanski et al., 2009; Fishman and Steinschneider, 2010). The one-dimensional CSD profile was generated from the local-field potentials and represents the laminar distribution of net transmembrane extracellular current flow associated with the synaptic activity of neural ensembles (Freeman and Nicholson, 1975; Nicholson and Freeman, 1975; Muller-Preuss and Mitzdorf, 1984; Mitzdorf, 1985; Steinschneider et al., 1992).

Our CSD profiles displayed laminar patterns of net current sources and sinks co-located with multi-unit activity that were typical of A1 in awake macaques (Steinschneider et al., 1992; Fishman et al., 2001a; Lakatos et al., 2005; Fishman and Steinschneider, 2010; Banno et al., 2023; Cai et al., 2025). An example CSD profile is shown in Fig. 1C, D. This profile had a dipole pattern indicating net current influx (a putative current sink) and efflux (a putative current source) that is characteristic of A1. In the middle electrode contacts, the CSD showed a sharp negative deflection (current sink) soon after stimulus onset. In the same contact, we also observed large increases in MUA (Fig. 1B). This large initial current sink, which typically coincided with the largest increases in MUA, is a characteristic feature of stimulus-evoked laminar response profiles in A1 and is consistent with post-synaptic depolarization of neural populations within putative input (granular) layers of the core auditory cortex (lamina 4 and lower lamina 3) (Steinschneider et al., 1992; Fishman et al., 2001b; Lakatos et al., 2005).

**Figure 1:**
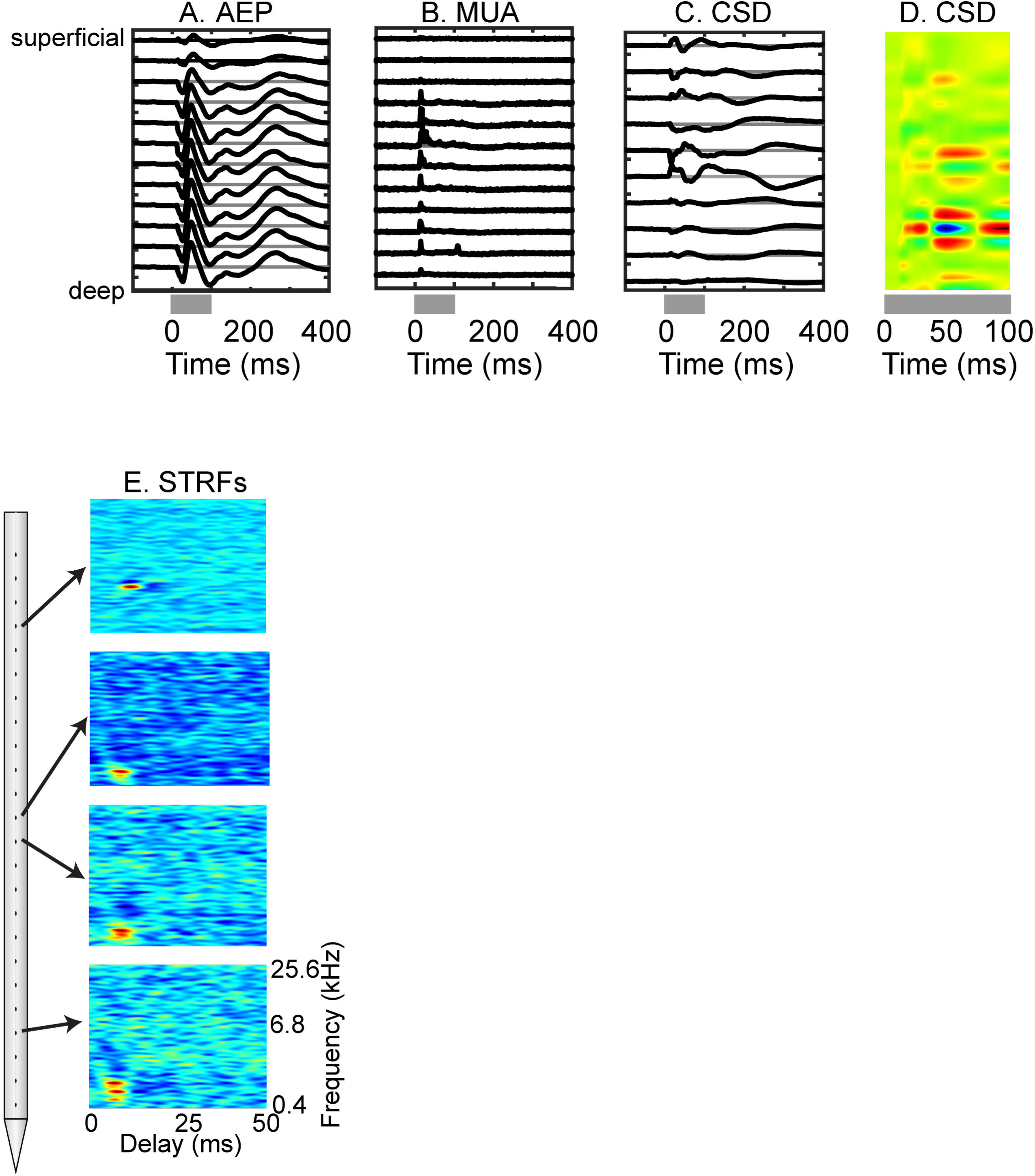
Representative laminar response profile and STRFs from A1. As a function of cortical depth from the superficial (top) to the deep (bottom) layers, we plot **(A)** the auditory evoked potentials (AEPs), which are derived from local-field potentials, **(B)** multiunit activity (MUA), and **(C)** the one-dimensional current source density (CSD), which is derived from the AEP profile. **(D)** In this color representation of the CSD, current sinks (net inward transmembrane current) are shown in blue, whereas current sources (net outward transmembrane current) are shown in red. The spacing between the electrode contacts is 200 µm. The amplitudes of these signals are in arbitrary units. The duration of the noise-burst stimulus (100 ms), which elicited the laminar response profile, is indicated by the grey bar under each panel. **(E)** Significant single-unit STRFs obtained from different electrode contacts during the same recording session. Arrows indicate the locations of the electrode contacts that had significant STRFs. The x-axis is aligned relative to stimulus onset. Color indicates the probability of eliciting a spike at each time-frequency point (red: maximum probability of eliciting a spike; blue: minimum probability of eliciting a spike).

We also found that spectrotemporal receptive fields (STRFs) showed features that are typical of A1. Example STRFs from electrode contacts along a single penetration are shown in Fig. 1E. Similar to our previous reports and others (Miller et al., 2002; Fritz et al., 2003; Atencio and Schreiner, 2010; Cai et al., 2025), our STRFs had short latencies with small circumscribed excitatory and/or inhibitory response fields. Further, we found that the best frequency (BF) of these STRFs was relatively constant along the electrode penetration (Cai et al., 2025). Our analyses reflect spiking activity obtained from all electrode contacts along the entire laminar extent of A1.

Consistent with previous studies that used the oddball paradigm (Fig. 2) in A1 (Ulanovsky et al., 2003; Farley et al., 2010; Fishman and Steinschneider, 2012; Natan et al., 2015; Parras et al., 2017), we found that the context of presentation modulated the neuron’s response to the BF tone burst. Specifically, we found that the firing rate was substantially higher to the BF tone burst when it was presented as the oddball (i.e., deviant stimulus) than when it was the standard stimulus. Single-neuron examples are shown in Figs. 3A and 4A. On average, we found that deviant stimuli elicited significantly higher A1 firing rates than standard stimuli (H_0_: median index value=0; the index is the difference between the deviant and standard firing rates, divided by their sum; Wilcoxon signed-rank test; Monkey W: N=61 single units; median [interquartile range, IQR]: 0.1 [−0.03-0.25]; p<0.002; Monkey C: N=38 single units; median [IQR]: 0.12 [−0.01-0.17]; p<0.001).

**Figure 2:**
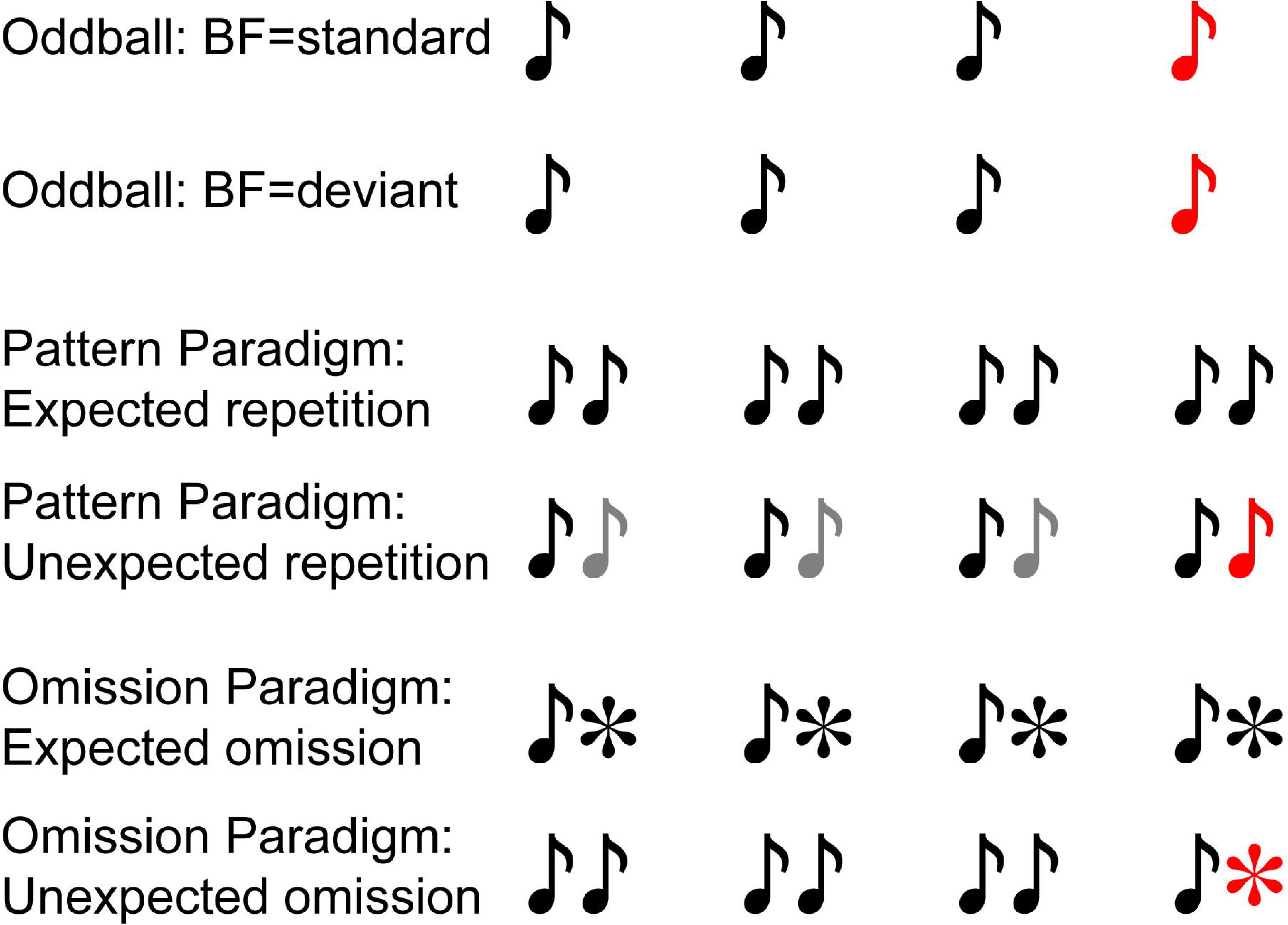
Schematic representation of the three auditory paradigms. Tone bursts are represented as “notes” with standard tone bursts represented in black and unexpected tone bursts in red. For the omission paradigm, the asterisk represents the expected (black) or the unexpected (red) missing tone burst.

**Figure 3:**
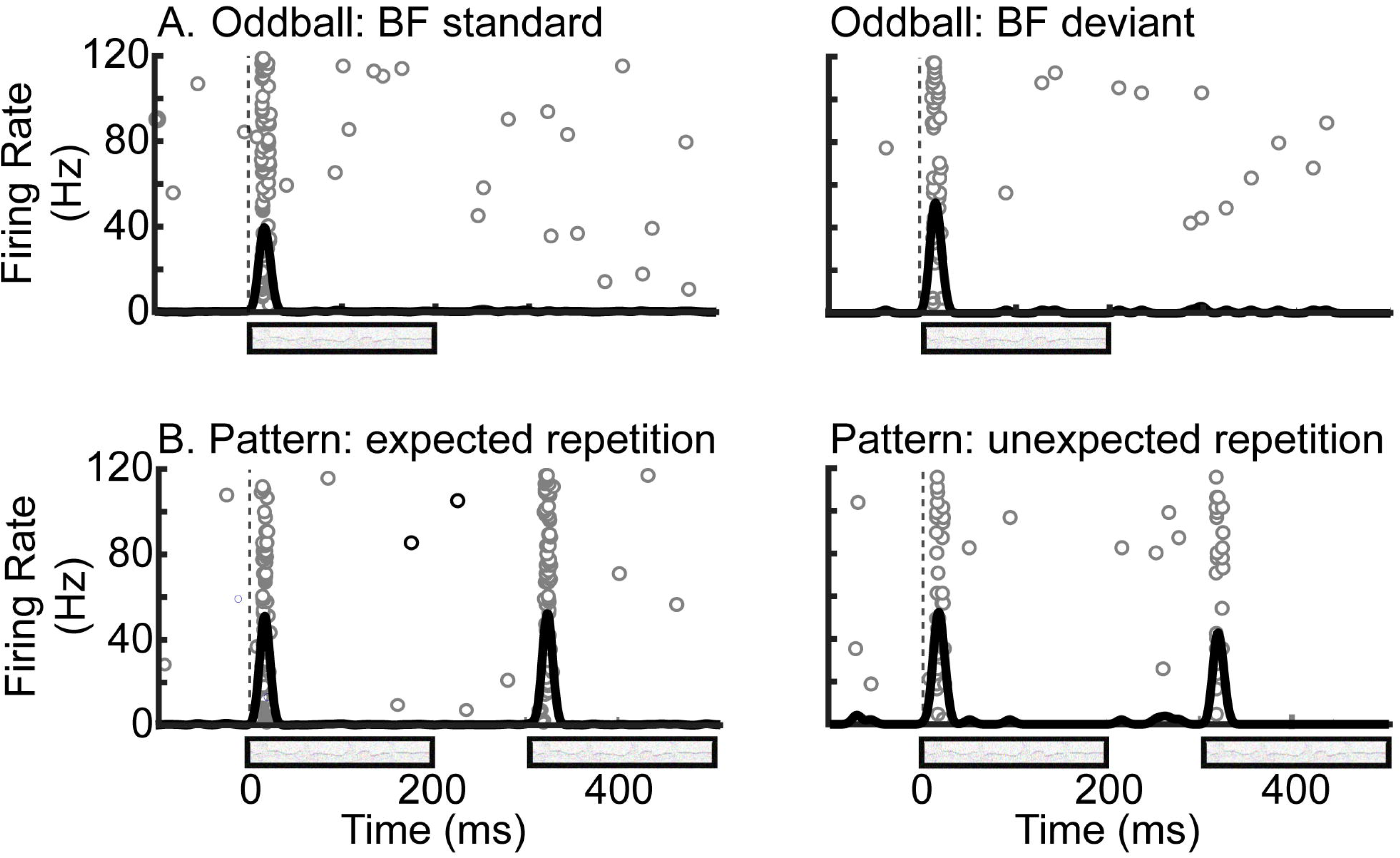
Raster and peristimulus-time histogram from a single neuron recorded during the oddball and pattern paradigms. **(A)** For the oddball paradigm, the raster plots and histograms are aligned relative to the onset of each tone burst. **(B)** For the pattern paradigm, the raster plots are aligned relative to the onset of each tone-burst pair. In both panels, the data in the left and right columns are data from the expected and unexpected conditions, respectively. The grey bars indicate the time course of a tone burst.

**Figure 4:**
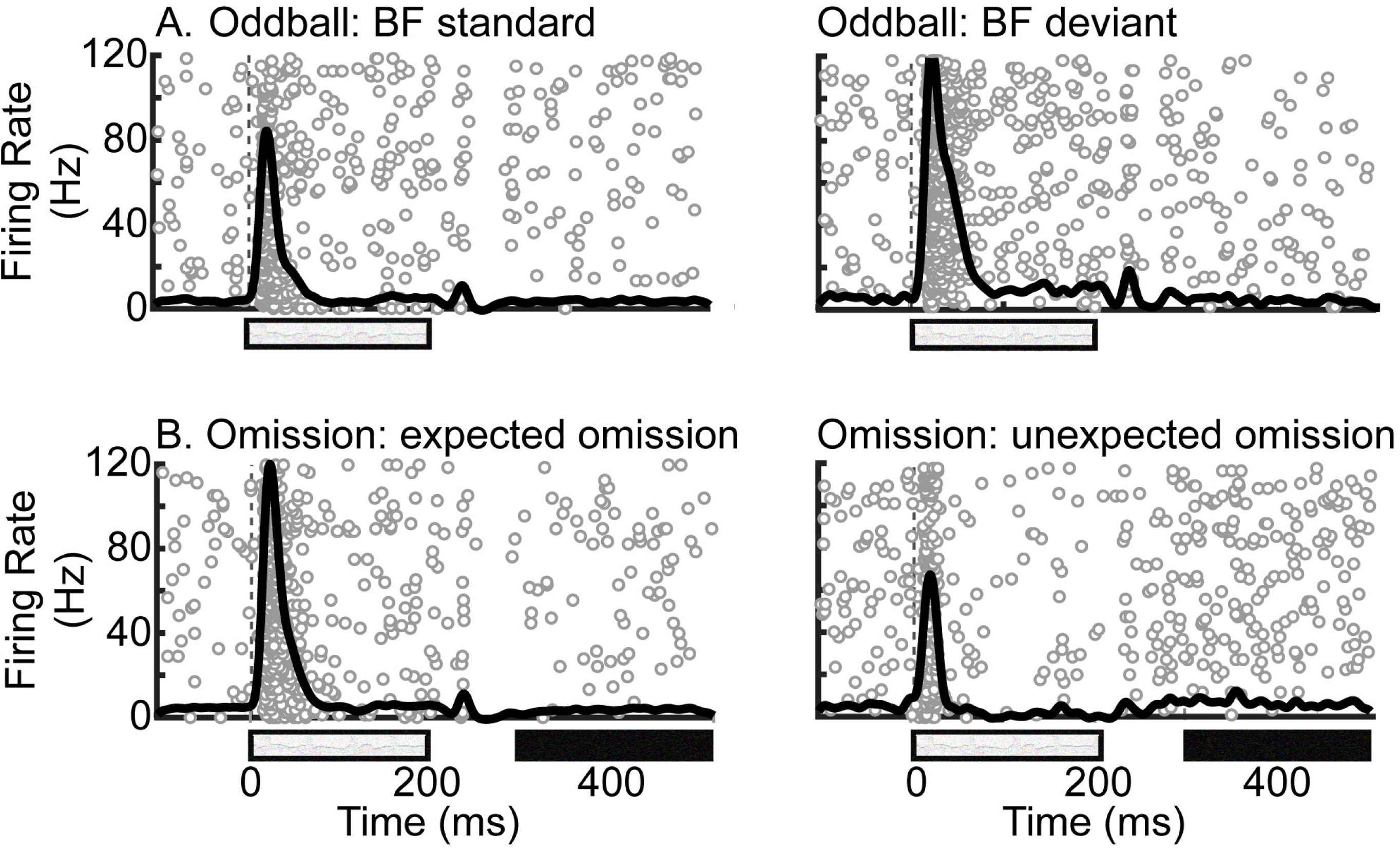
Raster and peristimulus-time histogram from a single neuron recorded during the oddball and omission paradigms. **(A)** For the oddball paradigm, the raster plots and histograms are aligned relative to the onset of each tone burst. **(B)** For the omission paradigm, the raster plots are aligned relative to the onset of the tone-burst + omitted-tone-burst complex. In both panels, the data in the left and right columns are data from the expected and unexpected conditions, respectively. The grey bars indicate the time course of a tone burst, whereas the black bars indicate the “time course” of the missing tone burst.

However, this pattern of responsivity is consistent with either reduced local suppression (e.g., due to SSA) or a prediction-error response as hypothesized by PC theory. To explore this issue further and to test these competing hypotheses, we probed A1 neurons with the pattern and omission paradigms (Fig. 2). Figs. 3-5 show the spiking patterns of three representative neurons recorded during the oddball, pattern, or admission paradigms.

**Figure 5:**
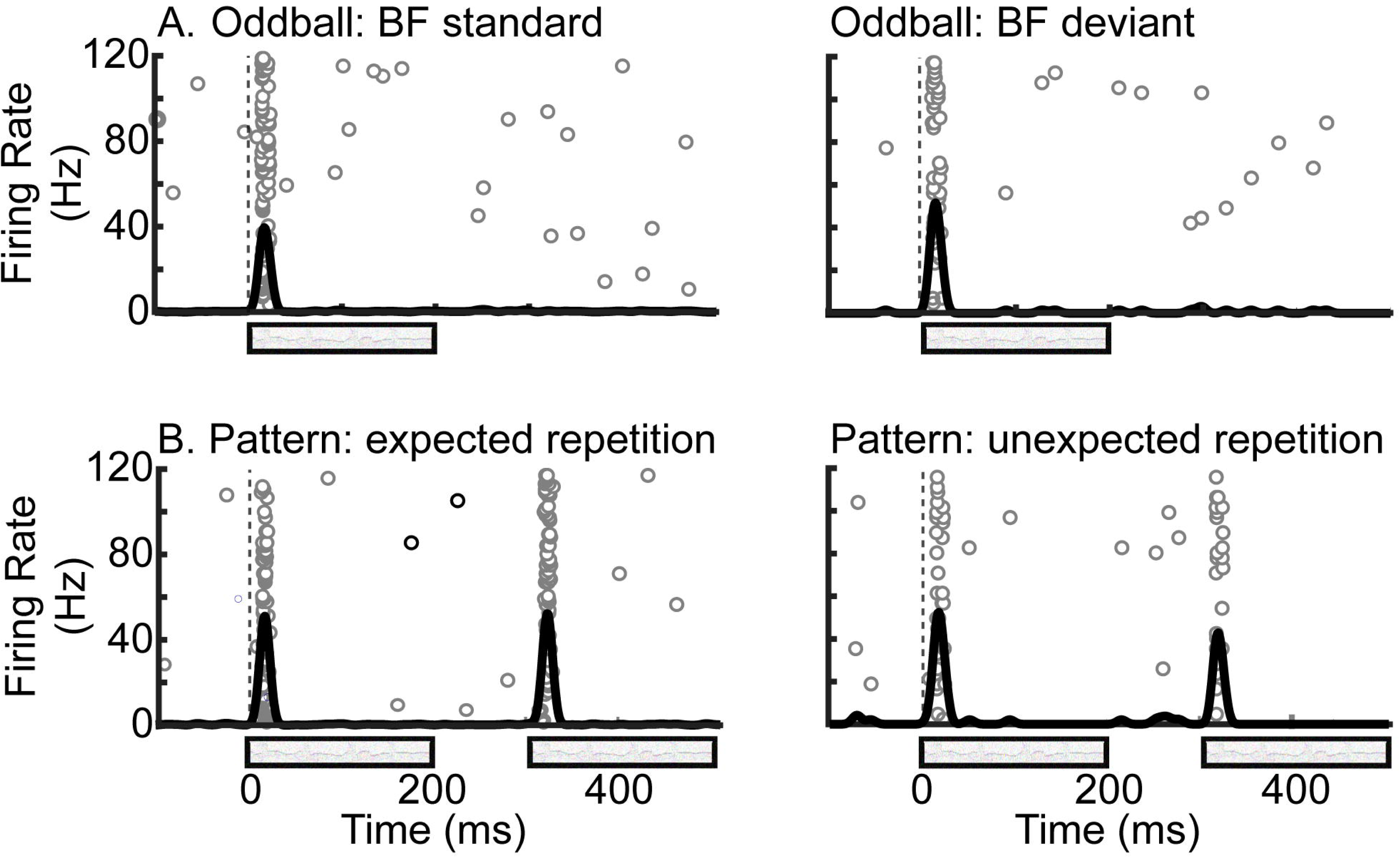
Raster and peristimulus-time histogram from a single neuron recorded during the pattern and omission paradigms. **(A)** For the pattern paradigm, the raster plots are aligned relative to the onset of each tone-burst pair. **(B)** For the omission paradigm, the raster plots are aligned relative to the onset of each tone-burst pair or tone-burst + omitted tone-burst complex. In both panels, the data in the left and right columns are data from the expected and unexpected conditions, respectively. The grey bars indicate the time course of a tone burst, whereas the black bars indicate the “time course” of the missing tone burst.

During the pattern paradigm, A1 neurons responded robustly and equally well to both expected versus unexpected tone-burst repetitions (Figs. 3B and 5A). That is, the unexpected tone-burst repetition (which is a violation of the expectation for alternation; see Fig. 2) did not substantially modulate the neuron’s firing rate. This finding is inconsistent with PC theory, which would predict a markedly larger response to the unexpected tone-burst repetition and would indicate the detection of a prediction error. In contrast, panels Figs. 4B and 5B show example response patterns during the omission paradigm. For these neurons, we found slight increases in firing rate in response to the unexpected tone-burst omission relative to the expected tone-burst omission, consistent with PC theory.

The response patterns of these single-neuron examples are representative of our population findings (Figs. 6 and 7). To quantify these observations, we calculated a prediction-error index (PEI; see **Materials and Methods**). PEI values ranged between −1 to +1. Values >0 indicate an enhanced response to the expectancy violation, consistent with PC, whereas values ≤0 indicate an altered or reduced response to the expectancy violation, inconsistent with PC.

**Figure 6:**
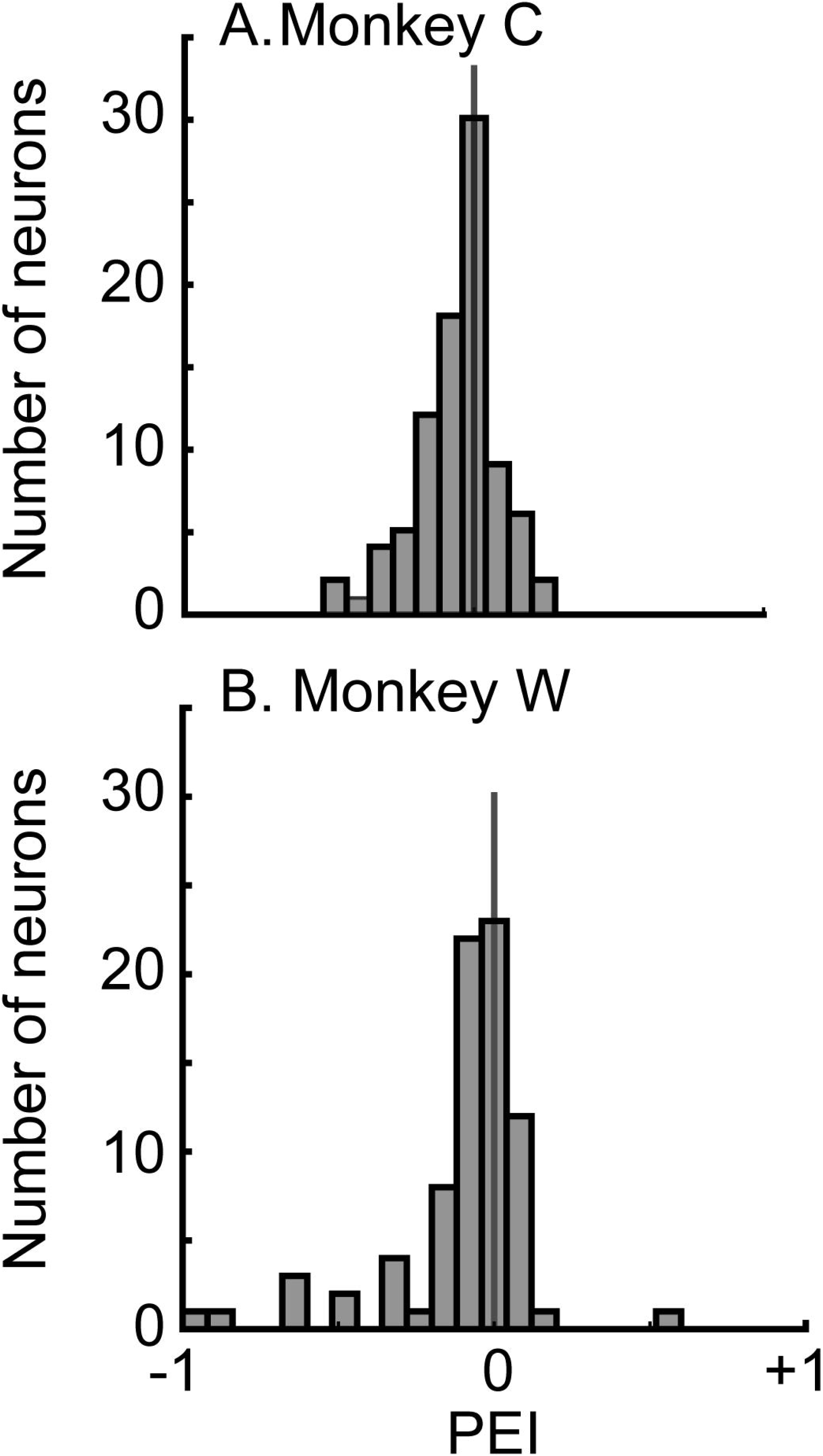
Distribution of PEI values during the pattern paradigm. PEI values >0 are consistent with predictive coding. The solid vertical line in each histogram indicates PEI=0. The data in **(A)** are from monkey C, whereas the data in **(B)** are from monkey W.

**Figure 7:**
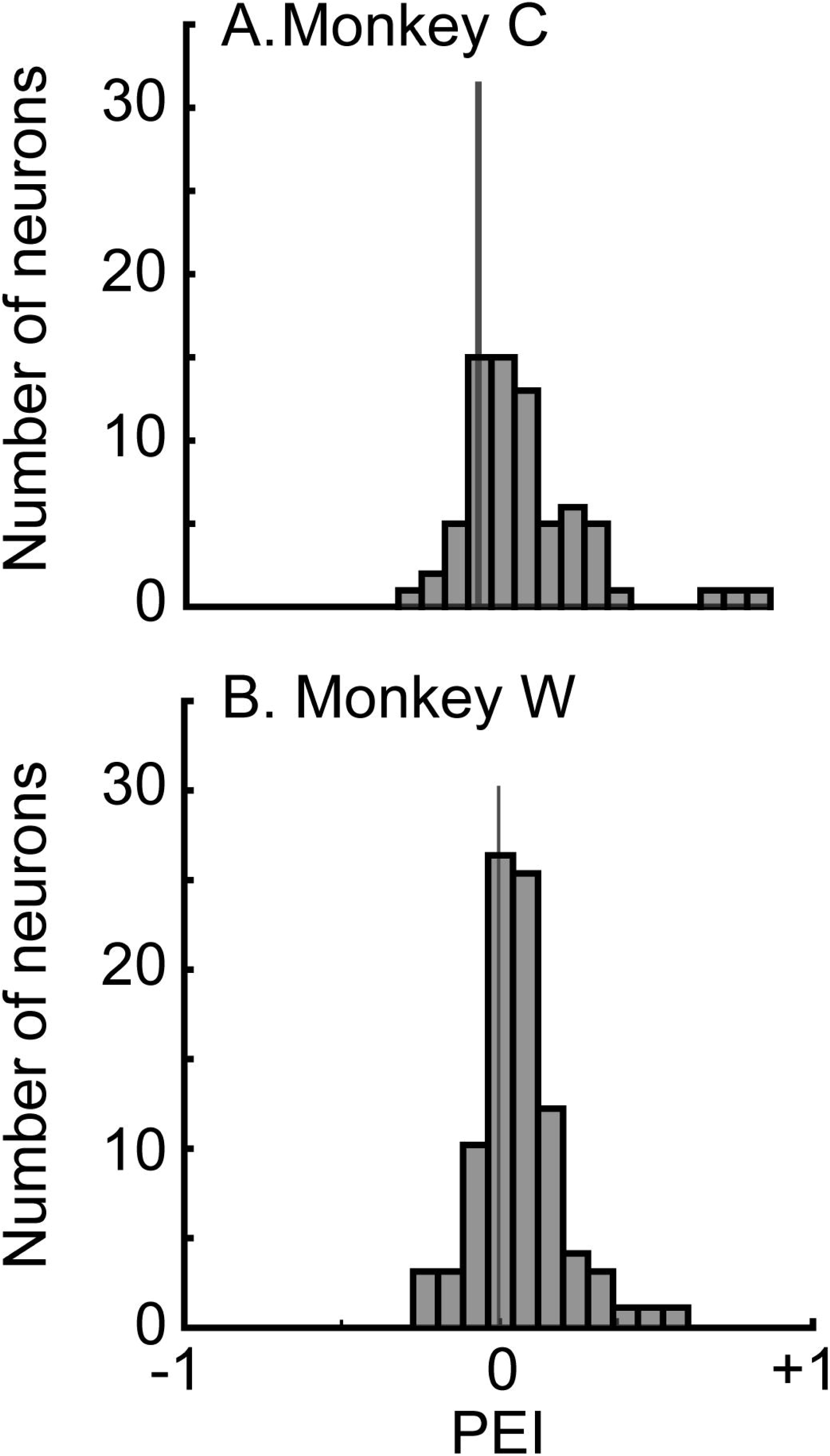
Distribution of PEI values from during the omission paradigm. PEI values >0 are consistent with predictive coding. The solid vertical line in each histogram indicates PEI=0. The data in **(A)** are from monkey C, whereas the data in **(B)** are from monkey W.

For the pattern paradigm (Fig. 6), the median PEI value was less than zero (H_0_: median index value=0; Wilcoxon signed-rank test; Monkey W: N=79; −0.05 [−0.12-0.02]; p<0.00001; Monkey C: N=89; −0.04 [−0.13-0.03]; p<0.0011), which is inconsistent with PC theory. In contrast, for the omission paradigm (Fig. 7), the median PEI value was greater than zero (H_0_: median index value=0; Wilcoxon signed-rank test; Monkey W: N=89; 0.05 [−0.004-0.12]; p<0.0001; Monkey C: N=71; 0.1 [0.01-0.24]; p<0.0001), consistent with PC theory.

Strengthening these conclusions further, a neuron-by-neuron analysis indicated that, on average, PEI values during the omission paradigm were larger than those during the pattern violation paradigm. This finding is shown in Fig. 8, where PEI values obtained for responses of individual neurons elicited during the omission paradigm are plotted against those elicited during the pattern paradigm. As can be seen, the majority of points lie below the line of equality, indicating that, on average, omission PEI values were greater than pattern PEI values (H_0_: median difference between the paired omission and pattern PEI values is zero; Monkey W: N=29; 0.06 [−0.053 – 0.15]; p<0.03; Monkey C: N=52 units that were recorded in both conditions; 0.13 [0.0001-0.29]; p<0.00002).

**Figure 8:**
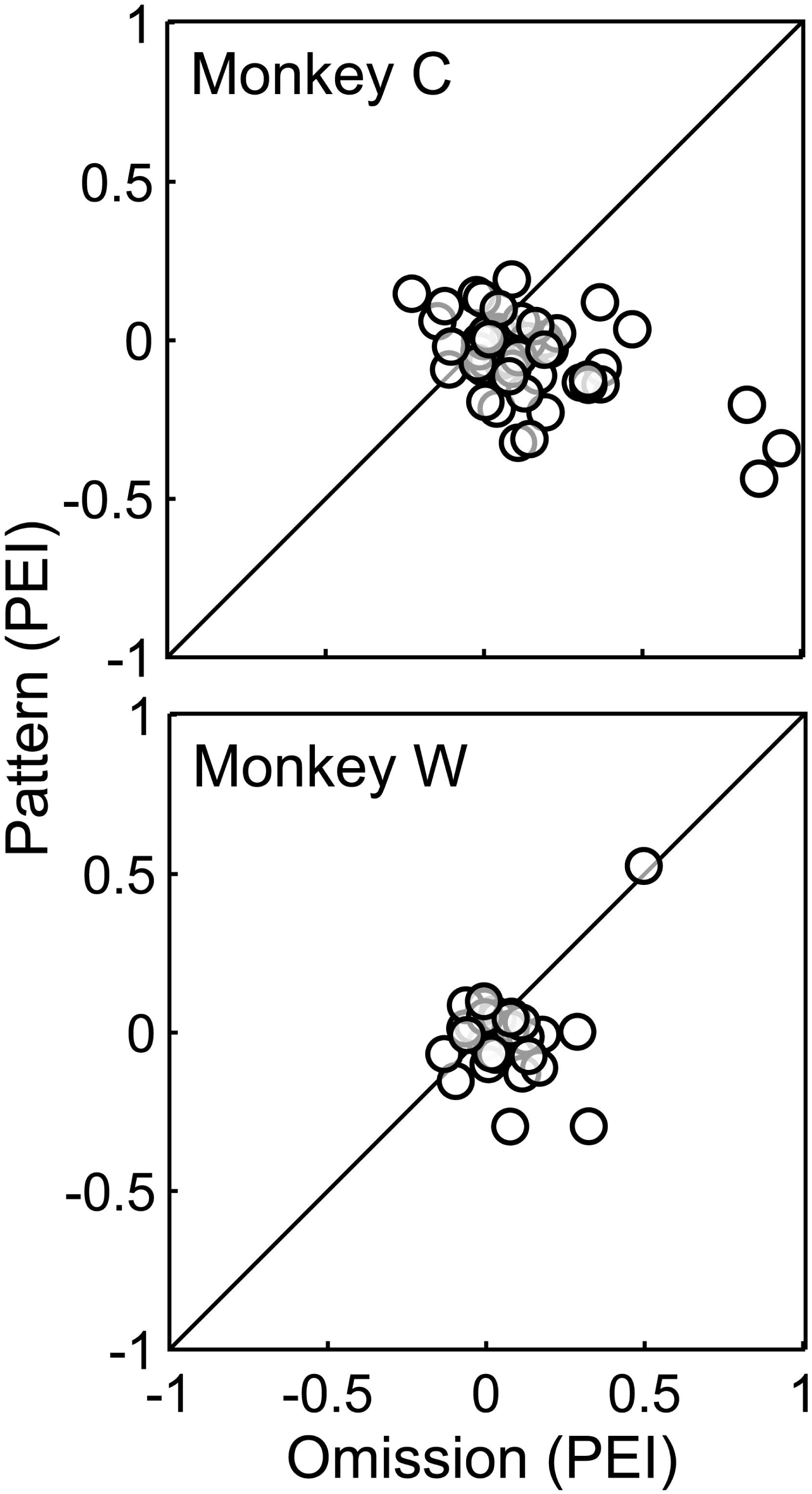
Correlation analysis of PEI values. On a neuron-by-neuron basis, the PEI values from the omission paradigm are plotted on the x-axis and the PEI values from the pattern paradigm are plotted on the y-axis. The data in **(A)** are from monkey C, whereas the data in **(B)** are from monkey W. The solid line in both panels is the line of unity.

Overall, our results indicate that the firing rates of individual A1 neurons are not enhanced for unexpected pattern violations and hence do not represent prediction errors. These findings are inconsistent with predictions of PC theory, but consistent with earlier preliminary findings in macaque A1 showing a similar absence of evidence for PC in neural population responses to unexpected stimulus repetitions (Fishman, 2014). On the other hand, the firing rates of individual A1 neurons are enhanced during unexpected stimulus omissions, thus providing evidential support for PC theory.

## Discussion

We tested whether there is evidence for the predictive-coding (PC) hypothesis in the primary auditory cortex (A1) of awake non-human primates. To isolate evidence for PC while controlling for SSA, we employed two auditory paradigms featuring unexpected pattern violations and unexpected stimulus omissions (Fishman, 2014). On average, A1 neurons were not modulated by expected versus unexpected stimulus repetitions (pattern violations), which is inconsistent with the predictions of PC. Conversely, and in line with PC, we found that A1 firing rates were higher for unexpected stimulus omissions compared to expected ones. Together, our findings provide nuanced evidence both for and against PC theory in A1.

Our results corroborate a substantial body of literature that has failed to find robust evidence for PC theory within A1, echoing previous findings of weak or absent prediction-error encoding during auditory-pattern violations. (Farley et al., 2010; Fishman and Steinschneider, 2010; Fishman, 2014; Parras et al., 2017; Polterovich et al., 2018). That is, SSA and other lower-level phenomena, such as forward suppression, appear to play a predominant role in modulating responses to temporally structured stimulus patterns. Of course, our null result does not preclude evidence for PC in non-primary areas of auditory cortex, as reported in previous human and non-human primate studies (Wacongne et al., 2011; Todorovic and de Lange, 2012; Uhrig et al., 2014; Dürschmid et al., 2016; Parras et al., 2017; Carbajal and Malmierca, 2018; Nourski et al., 2018; Hofmann-Shen et al., 2020). Nonetheless, insofar as primary sensory cortical areas are thought to be involved in the neural circuitry underlying PC, our findings provide important constraints on the neurobiological implementation of the theory.

Furthermore, our finding of enhanced A1 responses to unexpected stimulus omissions supports previous findings in other animal models and in humans (Hughes et al., 2001; Wacongne et al., 2011; Auksztulewicz et al., 2023; Cho et al., 2023; Lao-Rodríguez et al., 2023; Lao-Rodríguez et al., 2025; Yaron et al., 2025). Whereas our current data cannot definitively map the circuitry underlying omission responses, it likely involves top-down signals from higher-order regions that modulation sensory processing via feedback connections (Felleman and Van Essen, 1991; Shipp, 2007; Bastos et al., 2012; Markov and Kennedy, 2013; Chennu et al., 2016; Keller and Mrsic-Flogel, 2018; Bastos et al., 2020; Obara et al., 2023; Lao-Rodríguez et al., 2025).

These enhanced responses could also reflect signatures of the top-down predictions (expectations) themselves and not prediction errors per se (SanMiguel et al., 2013b; Kok et al., 2014; Schröger et al., 2015; Chennu et al., 2016; de Lange et al., 2018; Heilbron and Chait, 2018; Demarchi et al., 2019). Importantly, these putative prediction signals, although consistent with PC, do not uniquely support PC theory, as responses to stimulus omissions may be related to other processes (Halpern and Zatorre, 1999; Zatorre and Halpern, 2005; Chennu et al., 2016; Marion et al., 2021). Another possibility is that omission responses reflect involuntary attentional mechanisms that prioritize the unexpected absence of stimuli for enhanced processing (Salmi et al., 2009; Noyce et al., 2023; Baragona et al., 2025; Lao-Rodríguez et al., 2025). More generally, different variants of PC have been proposed, and PC is only one of several possible neural models of Bayesian perceptual inference in the brain (Aitchison and Lengyel, 2017; Spratling, 2017; Heilbron and Chait, 2018; Keller and Mrsic-Flogel, 2018; Sohn and Narain, 2021; Lange et al., 2023).

Several potential limitations of the present study should be noted. First, because we focused our analyses on responses to BF tone bursts, it is possible that stronger evidence for PC during the pattern violation paradigm may be reflected in non-BF responses. Second, we might have found stronger evidence for PC if the monkeys were actively engaged in a task as opposed to passive listening (Fritz et al., 2003; Niwa et al., 2012; Atiani et al., 2014; Downer et al., 2015; Bagur et al., 2018; Tsunada et al., 2019; Chillale et al., 2023). Third, more robust signatures of PC may not be evident in single-unit spiking activity but may be revealed in population-level activity (Fishman, 2014; Auksztulewicz et al., 2023). Finally, it remains an open question to what extent our findings obtained in macaques relate to neural substrates of predictive processing, statistical learning, and perceptual inference in humans.

## Materials and Methods

The University of Pennsylvania Institutional Animal Care and Use Committee approved all the experimental protocols in accordance with National Institutes of Health guidelines. The authors were not blind to group allocation during the experiment and when we analyzed the data outcomes.

All surgical procedures were conducted under general anesthesia and using aseptic surgical techniques. Two adult male rhesus macaques (*Macaca mulatta*; monkey W [14 years old] or monkey C [14 years old]) participated in this study. The monkeys were group housed and were maintained under water-controlled conditions during training and recording periods.

Before surgery, we identified the stereotactic location of the core auditory cortex (A1) through anatomical MRI scans (Romanski and Goldman-Rakic, 2002; Cohen et al., 2004; Frey et al., 2004; Johnston et al., 2016; Christison-Lagay et al., 2017). Using these scans, we designed a titanium recording chamber (Rogue, Inc) that was oriented perpendicular to the cortical layers of A1 (Frey et al., 2004; Banno et al., 2023; Cai et al., 2025). During surgery, we placed the monkey in a frameless stereotaxic device, removed the cranium over A1, and placed and secured the titanium chamber on the skull with titanium bone screws. Further details regarding the surgical approach can be found in Johnston et al. (Johnston et al., 2016).

We functionally defined A1 through two different complementary approaches (Banno et al., 2023; Cai et al., 2025). First, because we oriented our electrode array orthogonal to A1’s cortical lamina, we identified A1 recording sites by their characteristic laminar response profile, including current-source density (CSD; the second spatial derivative, relative to electrode-contact separation, of the local-field potential) and the spatial pattern of multiunit activity across the electrode contacts in a given penetration (Fig. 2) (Freeman and Nicholson, 1975; Nicholson and Freeman, 1975; Fishman et al., 2001a; O’Connell et al., 2014; Banno et al., 2023). Second, the anatomical arrangement of best frequencies obtained in different recording sessions was consistent with the anteroposterior tonotopic gradient characteristic of A1.

### Experimental chamber

In each session, a monkey was seated in a primate chair. We placed a calibrated free-field speaker (MICRO [Gallo Acoustics]) 1 m in front of the monkey and at their eye level. Calibrated auditory stimuli were generated with Matlab (Mathworks), passed through a digital-to-analog converter (E22, Lynx Studio Technology Inc), and presented through the free-field speaker. These sessions took place in an RF-shielded room with sound-attenuating walls and echo-absorbing foam on the inner walls.

### Auditory stimuli and paradigms

We presented dynamic moving ripple noise to generate a neuron’s spectrotemporal response field. From this response field, we calculated the site’s best frequency (BF; see *Neural analyses* below). On a day-by-day basis, we incorporated the BF into the frequency values of the tone bursts comprising the oddball, pattern, and omission paradigms (Fig. 2). In the pattern and omission paradigms, we established statistical regularities and violated those expectations. These paradigms were designed specifically to control for lower-level neurophysiological phenomena such as stimulus-specific adaptation, which cannot be distinguished from genuine prediction-error responses utilizing the classic oddball paradigm. The “pattern” paradigm tests for prediction-error responses elicited by unexpected tone-burst repetitions. In contrast, the “omission” paradigm tests for prediction error or prediction-related responses elicited by unexpected tone-bust omissions (Wacongne et al., 2011; Schröger et al., 2015; Chennu et al., 2016). During recording sessions in which we utilized the oddball and the pattern or omission paradigms, the tone bursts had a 200-ms duration. During those sessions in which only the pattern and omission paradigms were presented, the tone bursts had a 100-ms duration to match previous studies (Fishman, 2014). Regardless, all tone bursts had 5-ms linear on/off ramps, were presented with an intra-pair stimulus-onset asynchrony of 100 ms, an inter-pair stimulus-onset asynchrony of 458 ms, and at 65-70 dB SPL.

#### Dynamic Moving Ripple (DMR) Noise

DMR noise is a continuous time-varying broadband noise stimulus that covers the frequency range between 0.1 and 32 kHz (10-min duration; 65-dB spectrum level per ⅓ octave; 96-kHz sampling rate; 24-bit resolution) (Escabí and Schreiner, 2002; Escabi et al., 2014). At any instant of time, the stimulus has a sinusoidal spectrum; the density of the spectral peaks is determined by the spectral modulation frequency (Ω = 0-4 cycles/octave). The peak-to-peak amplitude of the ripple is 30 dB. The stimulus also contains temporal modulations (F_m_ = 0-25 Hz). Both the spectral and temporal parameters vary randomly and dynamically; the maximum rates of change are 0.25 Hz and 1 Hz, respectively.

#### Oddball Paradigm

This paradigm consisted of the repeated presentation of a tone burst at the same frequency value with the random presentation of a second “oddball” tone burst at a different frequency value. In the standard pattern, the frequency of the standard was set to the BF, and the frequency of the oddball was set a value that elicited half of the firing rate elicited by the BF tone burst. In the “violation” version, the standard frequency set a value that elicited half of the firing rate elicited by the BF tone burst, whereas the oddball was set to the BF.

#### Pattern Paradigm

This paradigm used two different tone-burst sequences. The first is a temporal sequence of identical tone-burst pairs separated by silence, which generated an expectation for tone-burst repetition (the “standard” pattern paradigm]; Fig. 2) (Alain et al., 1994; Macdonald and Campbell, 2011; Wacongne et al., 2011; Wacongne et al., 2012; Paavilainen, 2013; Fishman, 2014). The second was a temporal sequence of tone-burst pairs that alternated in frequency, which generated an expectation for tone-burst alternation. This expectation was then violated by the rare and unexpected repetition of one of the tone bursts (the “violation” pattern paradigm). The frequency of the first tone burst in the pair was set to the BF of the recording site. The frequency of the second tone-burst was set to a frequency value that elicited a neural response that was approximately half of that elicited by the BF tone burst (Fishman and Steinschneider, 2012).

#### Omission Paradigm

This paradigm used two tone-burst sequences conceptually similar to those used in earlier studies of omission responses in humans (Hughes et al., 2001). The first sequence consisted of the same-frequency tone-burst pairs, but the second tone burst in the pair was omitted 100% of the time; this generated an expectation for omission (the “standard” omission paradigm; Fig. 2). In contrast, the second sequence consisted of identical tone-burst pairs separated by silence, which generated an expectation for tone-burst repetition. This expectation for repetition was then violated by the rare, unexpected omission of the second tone burst in the pair (the “violation” omission paradigm; Fig. 2). The frequencies of the tone bursts were set to the site’s BF.

### Recording methodology and strategy

During a recording session, we first penetrated the dura with a stainless-steel guide tube (508-µm inner diameter) and advanced a 24-contact v-probe with 100-µm inter-contact spacing (Plexon) through the guide tube and into the brain. Neural signals were amplified (PZ5, Tucker-Davis Technologies) and digitized at 24.4 kHz (RZ2, Tucker-Davis Technologies) for both online and offline analyses.

While advancing the electrode, we presented an auditory “search stimulus” (an 80-ms white-noise burst at 65 dB SPL). When auditory-evoked responses were detected, we calculated the CSD profile from the neural activity generated from ∼110 repetitions of the search stimulus. Based on the CSD and multiunit activity (MUA) profiles, we adjusted the v-probe depth so that the electrode contacts with the highest MUA rates (Fig. 1A-C) and largest initial short-latency current sink were positioned in the middle contacts of the probe. This sink corresponds to extracellular current flow associated with postsynaptic potentials in stellate and pyramidal neurons within the thalamorecipient zone (lamina four and lower lamina) (Freeman and Nicholson, 1975; Nicholson and Freeman, 1975). Once this target depth was identified, we retracted the electrode by 250–500 µm and allowed the tissue to stabilize for ≥50 minutes to reduce electrode drift.

Next, we recorded neural activity while the monkey passively listened to the DMR stimulus (Escabí and Schreiner, 2002; Miller et al., 2002). We used this neural activity to generate the spectrotemporal receptive fields (STRFs) from isolated single-unit spiking activity and calculate its BF (Theunissen and Doupe, 1998; Theunissen et al., 2000; Sen et al., 2001; Theunissen et al., 2001; Grace et al., 2003). Next, we collected neural activity while monkeys listened to tone bursts presented within the context of the oddball, pattern, and/or omission paradigms. We presented the deviant stimuli either at consistent intervals within the violation variants of the paradigms (see Fig. 2) or at random times; in either case, deviant stimuli were only presented on 10–20% of the trials. Finally, at the end of each session, we regenerated both the site’s laminar CSD profile and the single-unit STRFs. We only report data from sessions in which the CSD and the STRFs were stable over the duration of the recording session. We delivered rewards manually and randomly throughout the recording session to help to ensure that the monkey remained awake. Neural signals were sampled at 24 kHz, filtered (0.003-6 kHz; PZ5-128, −64, RZ2, and RS4, TDT Inc.), and stored for online and offline analysis.

### Neural analyses

#### Extraction of Neural Signals

<u>Local-field potentials</u> (LFPs) were extracted by first low-pass filtering the neural signal (300-Hz cut-off frequency; four-pole bidirectional Butterworth filter) and then resampling the filtered signals at 1 kHz (Ghazanfar et al., 2005; Ghazanfar et al., 2008; Chandrasekaran and Ghazanfar, 2009; Tsunada et al., 2011; Gifford et al., 2019; Chiang et al., 2020). <u>Multiunit activity</u> (MUA) was extracted by filtering the neural signal (0.5-3.0 kHz), full-wave rectifying it, and then low-pass filtering (<0.6 kHz) to extract the envelope of summed action-potential activity (Lakatos et al., 2005; Kayser et al., 2007; Steinschneider et al., 2008; Fishman and Steinschneider, 2009). After high-pass filtering (0.6-6.0 kHz) the neural signal, <u>single-unit spiking activity</u> was sorted using Kilosort3 (Pachitariu et al., 2023). We used standard metrics to ensure high-quality single-neuron isolation (Schmitzer-Torbert et al., 2005; Liu et al., 2014).

#### Generation of a Neuron’s Spectrotemporal Response Field (STRF) and Determination of Best Frequency (BF)

We derived each neuron’s STRF using the reverse-correlation method, which is the average spectrotemporal stimulus envelope immediately preceding each spike elicited by the DMR stimulus (Miller et al., 2001; Escabí and Schreiner, 2002). A STRF’s “best frequency” (BF) was the frequency value that consistently elicited the largest neural response (Hermes et al., 1981; Kim and Young, 1994; Theunissen and Doupe, 1998; Keller and Takahashi, 2000; Theunissen et al., 2000; DePireux et al., 2001; Sen et al., 2001; Theunissen et al., 2001; Grace et al., 2003).

We utilized a two-step procedure to identify each STRF’s reliable samples and its overall reliability (Escabi et al., 2014). To identify each STRF’s significant spectrotemporal samples, we generated a null model under the assumption that spiking activity occurs randomly with respect to the DMR. We calculated this null model by first randomizing the inter-spike intervals and then averaging the spectrotemporal envelope of the DMR relative to the time of each spike. This generated a “noise” STRF. We repeated this procedure 1000× to generate a distribution of spectrotemporal noise-STRF samples. If a STRF sample was outside of the 99.9% confidence interval (i.e., p<0.001) of its respective noise-STRF sample, we considered it “significant”; if not, the STRF sample was assigned a value of 0.

For any given STRF, ∼230 STRF samples (i.e., 288 temporal×800 spectral samples×0.001) were false positives. Although these samples exceeded our significance criterion, it did not fulfill our primary objective of identifying STRFs with reproducible and reliable auditory responses. To identify such STRFs, we computed a “reliability” index by first breaking the DMR stimulus into twenty 30-s long segments (Escabi et al., 2014). Next, we randomly selected two subsamples of the 20 segments, generated a STRF from each of these segments, and calculated their Pearson correlation coefficient. A coefficient value of ∼1 (0) indicated that a given STRF is reliable (not reliable). We repeated this procedure 500× to generate a distribution of correlation-coefficient values. Finally, we generated a null distribution of coefficient values using the same process but with randomized inter-spike intervals. A Mann-Whitney test evaluated the hypothesis that the actual and the null correlation-coefficient distributions had the same median values. If the null hypothesis was rejected (p<0.01), we considered a STRF to be significant.

#### Analysis of Spiking Activity elicited During the Predictive-Coding Paradigms

To characterize the effect of the violations on neural activity, on a neuron-by-neuron basis, we computed a “response” index (RESI): RESI=(NR_2_-NR_1_)/(NR_2_+NR_1_), where NR_1_ and NR_2_ represent the mean firing rate elicited by the first and second tone burst, respectively, in a given tone-burst pair (see Fig. 2). We calculated the firing rate over a window extending from the onset of the tone burst to 50 ms after tone-burst offset. For the omission paradigm, we calculated the omitted response over an analogous time window. Next, we calculated the mean firing rate across all tone-burst pairs in a sequence. We generated two RESI values for the two tone-burst pattern sequences (expected and unexpected repetitions) and for the two tone-burst omission sequences (expected and unexpected omissions). Positive and negative RESI values imply repetition enhancement and suppression (e.g., stimulus-specific adaptation), respectively.

Finally, to quantify whether firing rate represented prediction errors in accordance with predictive coding theory, we computed a prediction-error index (PEI), which represents the difference between the RESI value calculated for the violation condition (i.e., unexpected repetition and omission) and the standard condition (i.e., expected repetition and omission), respectively. PEI values ranged between −1 to +1. Values >0 indicate an enhanced response to the expectancy violation, consistent with predictive coding, whereas values ≤0 indicate a reduced response to the expectancy violation, inconsistent with predictive coding.

### Statistical analyses

We used nonparametric tests on all analyses and null hypotheses were rejected at p<0.05.

## Acknowledgments

The authors want to thank Dr. Corey Roach for his help with stimulus development and analysis.

## Author contributions

BS, YF, and YEC designed the research; BS and HS performed the research; BS, LG, YF, and YEC analyzed data; BS, LG, HS, YF, and YEC wrote the paper

## References

Aitchison L, Lengyel M (2017) With or without you: predictive coding and Bayesian inference in the brain. Current opinion in neurobiology 46:219–227.

Alain C, Woods DL, Ogawa KH (1994) Brain indices of automatic pattern processing. Neuroreport 6:140–144.

Atencio CA, Schreiner CE (2010) Columnar connectivity and laminar processing in cat primary auditory cortex. PloS one 3:e9521.

Atiani S, David SV, Elgueda D, Locastro M, Radtke-Schuller S, Shamma SA, Fritz JB (2014) Emergent selectivity for task-relevant stimuli in higher-order auditory cortex. Neuron 82:486–499.

Auksztulewicz R, Rajendran VG, Peng F, Schnupp JWH, Harper NS (2023) Omission responses in local field potentials in rat auditory cortex. BMC Biol 21:130.

Bagur S, Averseng M, Elgueda D, David S, Fritz J, Yin P, Shamma S, Boubenec Y, Ostojic S (2018) Go/No-Go task engagement enhances population representation of target stimuli in primary auditory cortex. Nature communications 9:2529.

Banno T, Shirley H, Fishman YI, Cohen YE (2023) Changes in neural readout of response magnitude during auditory streaming do not correlate with behavioral choice in the auditory cortex. Cell reports 42:113493.

Baragona V, Schröger E, Widmann A (2025) Salient, Unexpected Omissions of Sounds Can Involuntarily Distract Attention. Journal of cognitive neuroscience:1–16.

Bastos AM, Lundqvist M, Waite AS, Kopell N, Miller EK (2020) Layer and rhythm specificity for predictive routing. Proceedings of the National Academy of Sciences of the United States of America 117:31459–31469.

Bastos AM, Usrey WM, Adams RA, Mangun GR, Fries P, Friston KJ (2012) Canonical microcircuits for predictive coding. Neuron 76:695–711.

Bendixen A, Schroger E, Winkler I (2009) I heard that coming: event-related potential evidence for stimulus-driven prediction in the auditory system. The Journal of neuroscience : the official journal of the Society for Neuroscience 29:8447–8451.

Cai H, Shirley H, Escabí MA, Cohen YE (2025) Distinct Cortical Populations Drive Multisensory Modulation of Segregated Auditory Sources. The Journal of neuroscience : the official journal of the Society for Neuroscience 45.

Carbajal GV, Malmierca MS (2018) The Neuronal Basis of Predictive Coding Along the Auditory Pathway: From the Subcortical Roots to Cortical Deviance Detection. Trends Hear 22:2331216518784822.

Chandrasekaran C, Ghazanfar AA (2009) Different neural frequency bands integrate faces and voices differently in the superior temporal sulcus. Journal of neurophysiology 101:773–788.

Chennu S, Noreika V, Gueorguiev D, Shtyrov Y, Bekinschtein TA, Henson R (2016) Silent Expectations: Dynamic Causal Modeling of Cortical Prediction and Attention to Sounds That Weren’t. The Journal of neuroscience : the official journal of the Society for Neuroscience 36:8305–8316.

Chiang CH, Lee J, Wang C, Williams AJ, Lucas TH, Cohen YE, Viventi J (2020) A modular high-density µECoG system on macaque vlPFC for auditory cognitive decoding. Journal of neural engineering 17:046008.

Chillale RK, Shamma S, Ostojic S, Boubenec Y (2023) Dynamics and maintenance of categorical responses in primary auditory cortex during task engagement. eLife 12.

Cho H, Fonken YM, Adamek M, Jimenez R, Lin JJ, Schalk G, Knight RT, Brunner P (2023) Unexpected sound omissions are signaled in human posterior superior temporal gyrus: an intracranial study. Cerebral cortex (New York, NY : 1991) 33:8837–8848.

Christison-Lagay KL, Bennur S, Cohen YE (2017) Contribution of spiking activity in the primary auditory cortex to detection in noise. J Neurophysiolology 118:3118–3131.

Cohen YE, Cohen IS, Gifford III GW (2004) Modulation of LIP activity by predictive auditory and visual cues. Cerebral Cortex 14:1287–1301.

de Lange FP, Heilbron M, Kok P (2018) How Do Expectations Shape Perception? Trends in cognitive sciences 22:764–779.

Demarchi G, Sanchez G, Weisz N (2019) Automatic and feature-specific prediction-related neural activity in the human auditory system. Nature communications 10:3440.

Denham SL, Winkler I (2006) The role of predictive models in the formation of auditory streams. Journal of physiology, Paris 100:154–170.

DePireux DA, Simon JZ, Klein DJ, Shamma SA (2001) Spectro-temporal response field characterization with dynamic ripples in ferret primary auditory cortex. Journal of neurophysiology 85:1220–1234.

Downer JD, Niwa M, Sutter ML (2015) Task engagement selectively modulates neural correlations in primary auditory cortex. Journal Neuroscience 35:7565–7574.

Dürschmid S, Edwards E, Reichert C, Dewar C, Hinrichs H, Heinze HJ, Kirsch HE, Dalal SS, Deouell LY, Knight RT (2016) Hierarchy of prediction errors for auditory events in human temporal and frontal cortex. Proceedings of the National Academy of Sciences of the United States of America 113:6755–6760.

Escabi MA, Read HL, Viventi J, Kim DH, Higgins NC, Storace DA, Liu AS, Gifford AM, Burke JF, Campisi M, Kim YS, Avrin AE, Spiegel Jan V, Huang Y, Li M, Wu J, Rogers JA, Litt B, Cohen YE (2014) A high-density, high-channel count, multiplexed muECoG array for auditory-cortex recordings. Journal of neurophysiology 112:1566–1583.

Escabí MA, Schreiner CE (2002) Nonlinear spectrotemporal sound analysis by neurons in the auditory midbrain. The Journal of neuroscience : the official journal of the Society for Neuroscience 22:4114–4131.

Farley BJ, Quirk MC, Doherty JJ, Christian EP (2010) Stimulus-specific adaptation in auditory cortex is an NMDA-independent process distinct from the sensory novelty encoded by the mismatch negativity. The Journal of neuroscience : the official journal of the Society for Neuroscience 30:16475–16484.

Felleman DJ, Van Essen DC (1991) Distributed hierarchical processing in the primate cerebral cortex. Cerebral cortex (New York, NY : 1991) 1:1–47.

Fishman YI (2014) The mechanisms and meaning of the mismatch negativity. Brain topography 27:500–526.

Fishman YI, Steinschneider M (2009) Temporally dynamic frequency tuning of population responses in monkey primary auditory cortex. Hearing research 254:64–76.

Fishman YI, Steinschneider M (2010) Neural Correlates of Auditory Scene Analysis Based on Inharmonicity in Monkey Primary Auditory Cortex. Journal of Neuroscience 30:12480–12494.

Fishman YI, Steinschneider M (2012) Searching for the mismatch negativity in primary auditory cortex of the awake monkey: deviance detection or stimulus specific adaptation? The Journal of neuroscience : the official journal of the Society for Neuroscience 32:15747–15758.

Fishman YI, Reser DH, Arezzo JC, Steinschneider M (2001a) Neural correlates of auditory stream segregation in primary auditory cortex of the awake monkey. Hearing research 151:167–187.

Fishman YI, Volkov IO, Noh MD, Garell PC, Bakken H, Arezzo JC, Howard MA, Steinschneider M (2001b) Consonance and dissonance of musical chords: neural correlates in auditory cortex of monkeys and humans. J Neurophysiology 86:2761–2788.

Freeman JA, Nicholson C (1975) Experimental optimization of current source-density technique for anuran cerebellum. Journal of neurophysiology 38:369–382.

Frey S, Comeau R, Hynes B, Mackey S, Petrides M (2004) Frameless stereotaxy in the nonhuman primate. NeuroImage 23:1226–1234.

Friston K (2005) A theory of cortical responses. Philosophical transactions of the Royal Society of London Series B, Biological sciences 360:815–836.

Fritz J, Shamma S, Elhilali M, Klein D (2003) Rapid task-related plasticity of spectrotemporal receptive fields in primary auditory cortex. Nature neuroscience 6:1216–1223.

Ghazanfar AA, Chandrasekaran C, Logothetis NK (2008) Interactions between the superior temporal sulcus and auditory cortex mediate dynamic face/voice integration in rhesus monkeys. The Journal of neuroscience : the official journal of the Society for Neuroscience 28:4457–4469.

Ghazanfar AA, Maier JX, Hoffman KL, Logothetis NK (2005) Multisensory integration of dynamic faces and voices in rhesus monkey auditory cortex. The Journal of neuroscience : the official journal of the Society for Neuroscience 25:5004–5012.

Gifford AM, Sperling MR, Sharan A, Gorniak RJ, Williams RB, Davis K, Kahana MJ, Cohen YE (2019) Neuronal phase consistency tracks dynamic changes in acoustic spectral regularity. European Journal of Neuroscience 49:1268–1287.

Gold JI, Shadlen MN (2002) Banburismus and the brain: decoding the relationship beteween sensory stimuli, decisions, and reward. Neuron 36:299–308.

Gold JI, Shadlen MN (2007) The Neural Basis of Decision Making. Annual review of neuroscience 30:535–574.

Grace JA, Amin N, Singh NC, Theunissen FE (2003) Selectivity for conspecific song in the zebra finch auditory forebrain. Journal of neurophysiology 89:472–487.

Griffiths TL, Tenenbaum JB (2011) Predicting the future as Bayesian inference: people combine prior knowledge with observations when estimating duration and extent. J Exp Psychol Gen 140:725–743.

Halpern AR, Zatorre RJ (1999) When That Tune Runs Through Your Head: A PET Investigation of Auditory Imagery for Familiar Melodies. Cerebral cortex (New York, NY : 1991) 9:697–704.

Heilbron M, Chait M (2018) Great Expectations: Is there Evidence for Predictive Coding in Auditory Cortex? Neuroscience 389:54–73.

Hermes DJ, Aertsen AM, Johannesma PI, Eggermont JJ (1981) Spectrotemporal characteristics of single units in the auditory midbrain of the lightly anesthetised grass frog (*Rana temporaria* L.) investigated with tonal stimuli. Hearing research 6:103–126.

Hofmann-Shen C, Vogel BO, Kaffes M, Rudolph A, Brown EC, Tas C, Brüne M, Neuhaus AH (2020) Mapping adaptation, deviance detection, and prediction error in auditory processing. Neuroimage 207:116432.

Hughes HC, Darcey TM, Barkan HI, Williamson PD, Roberts DW, Aslin CH (2001) Responses of Human Auditory Association Cortex to the Omission of an Expected Acoustic Event. NeuroImage 13:1073–1089.

Johnston JM, Cohen YE, Shirley H, Tsunada J, Bennur S, Christison-Lagay K, Veeder CL (2016) Recent refinements to cranial implants for rhesus macaques (Macaca mulatta). Lab animal 45:180–186.

Kayser C, Petkov CI, Logothetis NK (2007) Tuning to sound frequency in auditory field potentials. Journal of neurophysiology 98:1806–1809.

Keller CH, Takahashi TT (2000) Representation of temporal features of complex sounds by the discharge patterns of neurons in the owl’s inferior colliculus. Journal of neurophysiology 84:2638–2650.

Keller GB, Mrsic-Flogel TD (2018) Predictive Processing: A Canonical Cortical Computation. Neuron 100:424–435.

Kim PJ, Young ED (1994) Comparative analysis of spectro-temporal receptive fields, reverse correlation functions, and frequency tuning curves of auditory-nerve fibers. The Journal of the Acoustical Society of America 95:410–422.

Knill DC, Richards W, eds (1996) Perception as (Bayesian) Inference. Cambridge, UK: Cambridge University Press.

Koelsch S, Vuust P, Friston K (2019) Predictive Processes and the Peculiar Case of Music. Trends in cognitive sciences 23:63–77.

Kok P, Failing MF, de Lange FP (2014) Prior expectations evoke stimulus templates in the primary visual cortex. Journal of cognitive neuroscience 26:1546–1554.

Lakatos P, Shah AS, Knuth KH, Ulbert I, Karmos G, Schroeder CE (2005) An oscillatory hierarchy controlling neuronal excitability and stimulus processing in the auditory cortex. Journal of neurophysiology 94:1904–1911.

Lange RD, Shivkumar S, Chattoraj A, Haefner RM (2023) Bayesian encoding and decoding as distinct perspectives on neural coding. Nature neuroscience 26:2063–2072.

Lao-Rodríguez AB, Schröger E, Malmierca MS (2025) The sound of silence: Omission responses and how the brain predicts in the absence of sound. Neuroscience and biobehavioral reviews 181:106505.

Lao-Rodríguez AB, Przewrocki K, Pérez-González D, Alishbayli A, Yilmaz E, Malmierca MS, Englitz B (2023) Neuronal responses to omitted tones in the auditory brain: A neuronal correlate for predictive coding. Sci Adv 9:eabq8657.

Liu X, Wan H, Shi L (2014) Quality metrics of spike sorting using neighborhood components analysis. The open biomedical engineering journal 8:60–67.

Macdonald M, Campbell K (2011) Effects of a violation of an expected increase or decrease in intensity on detection of change within an auditory pattern. Brain Cogn 77:438–445.

Marion G, Di Liberto GM, Shamma SA (2021) The Music of Silence: Part I: Responses to Musical Imagery Encode Melodic Expectations and Acoustics. The Journal of neuroscience : the official journal of the Society for Neuroscience 41:7435–7448.

Markov NT, Kennedy H (2013) The importance of being hierarchical. Current opinion in neurobiology 23:187–194.

May PJ, Tiitinen H (2010) Mismatch negativity (MMN), the deviance-elicited auditory deflection, explained. Psychophysiology 47:66–122.

May PJC (2021) The Adaptation Model Offers a Challenge for the Predictive Coding Account of Mismatch Negativity. Frontiers in human neuroscience 15.

Miller LM, Escabí MA, Read HL, Schreiner CE (2001) Feature selectivity and interneuronal cooperation in the thalamocortical system. The Journal of neuroscience : the official journal of the Society for Neuroscience 21:8136–8144.

Miller LM, Escabí MA, Read HL, Schreiner CE (2002) Spectrotemporal receptive fields in the lemniscal auditory thalamus and cortex. Journal of neurophysiology 87:516–527.

Mitzdorf U (1985) Current source-density method and application in cat cerebral cortex: investigation of evoked potentials and EEG phenomena. Physiological reviews 65:37–100.

Muller-Preuss P, Mitzdorf U (1984) Functional anatomy of the inferior colliculus and the auditory cortex: current source density analyses of click-evoked potentials. Hearing research 16:133–142.

Natan RG, Briguglio JJ, Mwilambwe-Tshilobo L, Jones SI, Aizenberg M, Goldberg EM, Geffen MN (2015) Complementary control of sensory adaptation by two types of cortical interneurons. eLife 4.

Nicholson C, Freeman JA (1975) Theory of current source-density analysis and determination of conductivity tensor for anuran cerebellum. Journal of neurophysiology 38:356–368.

Niwa M, Johnson JS, O’Connor KN, Sutter ML (2012) Active Engagement Improves Primary Auditory Cortical Neurons’ Ability to Discriminate Temporal Modulation. Journal of Neuroscience 32:9323–9334.

Nourski KV, Steinschneider M, Rhone AE, Kawasaki H, Howard MA, 3rd, Banks MI (2018) Auditory Predictive Coding across Awareness States under Anesthesia: An Intracranial Electrophysiology Study. The Journal of neuroscience : the official journal of the Society for Neuroscience 38:8441–8452.

Noyce AL, Kwasa JAC, Shinn-Cunningham BG (2023) Defining attention from an auditory perspective. Wiley Interdiscip Rev Cogn Sci 14:e1610.

O’Connell MN, Barczak A, Schroeder CE, Lakatos P (2014) Layer Specific Sharpening of Frequency Tuning by Selective Attention in Primary Auditory Cortex. The Journal of Neuroscience 34:16496–16508.

O’Reilly JA (2021) Roving oddball paradigm elicits sensory gating, frequency sensitivity, and long-latency response in common marmosets. IBRO Neurosci Rep 11:128–136.

Obara K, Ebina T, Terada SI, Uka T, Komatsu M, Takaji M, Watakabe A, Kobayashi K, Masamizu Y, Mizukami H, Yamamori T, Kasai K, Matsuzaki M (2023) Change detection in the primate auditory cortex through feedback of prediction error signals. Nature communications 14:6981.

Paavilainen P (2013) The mismatch-negativity (MMN) component of the auditory event-related potential to violations of abstract regularities: a review. International journal of psychophysiology : official journal of the International Organization of Psychophysiology 88:109–123.

Pachitariu M, Steinmetz N, Kadir S, Carandini M, Harris KD (2023) Kilosort: realtime spike-sorting for extracellular electrophysiology with hundreds of channels. bioRxiv:061481.

Parras GG, Nieto-Diego J, Carbajal GV, Valdés-Baizabal C, Escera C, Malmierca MS (2017) Neurons along the auditory pathway exhibit a hierarchical organization of prediction error. Nature communications 8:2148.

Polterovich A, Jankowski MM, Nelken I (2018) Deviance sensitivity in the auditory cortex of freely moving rats. PloS one 13:e0197678.

Rao RPN, Ballard DH (1999) Predictive coding in the visual cortex: a functional interpretation of some extra-classical receptive-field effects. Nature neuroscience 2:79–87.

Romanski LM, Goldman-Rakic PS (2002) An auditory domain in primate prefrontal cortex. Nature neuroscience 5:15–16.

Ross JM, Hamm JP (2020) Cortical Microcircuit Mechanisms of Mismatch Negativity and Its Underlying Subcomponents. Frontiers in neural circuits 14:13.

Salmi J, Rinne T, Koistinen S, Salonen O, Alho K (2009) Brain networks of bottom-up triggered and top-down controlled shifting of auditory attention. Brain research 1286:155–164.

Sanmiguel I, Saupe K, Schröger E (2013a) I know what is missing here: electrophysiological prediction error signals elicited by omissions of predicted “what” but not “when”. Frontiers in human neuroscience 7:407.

SanMiguel I, Widmann A, Bendixen A, Trujillo-Barreto N, Schröger E (2013b) Hearing silences: human auditory processing relies on preactivation of sound-specific brain activity patterns. The Journal of neuroscience : the official journal of the Society for Neuroscience 33:8633–8639.

Schmitzer-Torbert N, Jackson J, Henze D, Harris K, Redish AD (2005) Quantitative measures of cluster quality for use in extracellular recordings. Neuroscience 131:1–11.

Schröger E, Marzecová A, SanMiguel I (2015) Attention and prediction in human audition: a lesson from cognitive psychophysiology. The European journal of neuroscience 41:641–664.

Sen K, Theunissen FE, Doupe AJ (2001) Feature analysis of natural sounds in the songbird auditory forebrain. Journal of neurophysiology 85:1445–1458.

Shipp S (2007) Structure and function of the cerebral cortex. Current biology : CB 17:R443–R339.

Shipp S (2016) Neural Elements for Predictive Coding. Frontiers in psychology 7:1792.

Sohn H, Narain D (2021) Neural implementations of Bayesian inference. Current opinion in neurobiology 70:121–129.

Spratling MW (2017) A review of predictive coding algorithms. Brain Cogn 112:92–97.

Steinschneider M, Fishman YI, Arezzo J, C. (2008) Spectrotemporal analysis of evoked and induced electroencephalographic responses in primary auditory cortex (A1) of the awake monkey. Cerebral cortex (New York, NY : 1991) 18:610–625.

Steinschneider M, Tenke CE, Schroeder CE, Javitt DC, Simpson GV, Arezzo JC, Vaughan HG, Jr. (1992) Cellular generators of the cortical auditory evoked potential initial component. Electroencephalography and clinical neurophysiology 84:196–200.

Symonds RM, Lee WW, Kohn A, Schwartz O, Witkowski S, Sussman ES (2017) Distinguishing Neural Adaptation and Predictive Coding Hypotheses in Auditory Change Detection. Brain topography 30:136–148.

Szabó BT, Denham SL, Winkler I (2016) Computational Models of Auditory Scene Analysis: A Review. Frontiers in Neuroscience Volume 10 - 2016.

Szymanski FD, Garcia-Lazaro JA, Schnupp JW (2009) Current source density profiles of stimulus-specific adaptation in rat auditory cortex. Journal of neurophysiology 102:1483–1490.

Teichert T, Jedema H, Zhen Z, Gurnsey K (2025) Complementary functional profiles of mismatch responses mediated by adaptation and deviance detection point to two distinct auditory short-term memory systems. Journal of neurophysiology 134:314–336.

Theunissen FE, Doupe AJ (1998) Temporal and spectral sensitivity of complex auditory neurons in the nucleus HVc of male zebra finches. The Journal of neuroscience : the official journal of the Society for Neuroscience 18:3786–3802.

Theunissen FE, Sen K, Doupe AJ (2000) Spectral-temporal receptive fields of nonlinear auditory neurons obtained using natural sounds. The Journal of neuroscience : the official journal of the Society for Neuroscience 20:2315–2331.

Theunissen FE, David SV, Singh NC, Hsu A, Vinje WE, Gallant JL (2001) Estimating spatio-temporal receptive fields of auditory and visual neurons from their responses to natural stimuli. Network 12:289–316.

Todorovic A, de Lange FP (2012) Repetition suppression and expectation suppression are dissociable in time in early auditory evoked fields. The Journal of neuroscience : the official journal of the Society for Neuroscience 32:13389–13395.

Tsunada J, Cohen Y, Gold JI (2019) Post-decision processing in primate prefrontal cortex influences subsequent choices on an auditory decision-making task. eLife 8:e46770.

Tsunada J, Baker AE, Christison-Lagay KL, Davis SJ, Cohen YE (2011) Modulation of cross-frequency coupling by novel and repeated stimuli in the primate ventrolateral prefrontal cortex. Front Psychology 2.

Uhrig L, Dehaene S, Jarraya B (2014) A hierarchy of responses to auditory regularities in the macaque brain. The Journal of neuroscience : the official journal of the Society for Neuroscience 34:1127–1132.

Ulanovsky N, Las L, Nelken I (2003) Processing of low-probability sounds by cortical neurons. Nature neuroscience 6:391–398.

Wacongne C, Changeux JP, Dehaene S (2012) A neuronal model of predictive coding accounting for the mismatch negativity. The Journal of neuroscience : the official journal of the Society for Neuroscience 32:3665–3678.

Wacongne C, Labyt E, van Wassenhove V, Bekinschtein T, Naccache L, Dehaene S (2011) Evidence for a hierarchy of predictions and prediction errors in human cortex. Proceedings of the National Academy of Sciences of the United States of America 108:20754–20759.

Yaron A, Shiramatsu-Isoguchi T, Kern FB, Ohki K, Takahashi H, Chao ZC (2025) Auditory cortex neurons that encode negative prediction errors respond to omissions of sounds in a predictable sequence. PLoS biology 23:e3003242.

Zarcone A, van Schijndel M, Vogels J, Demberg V (2016) Salience and Attention in Surprisal-Based Accounts of Language Processing. Frontiers in psychology Volume 7–2016.

Zatorre RJ, Halpern AR (2005) Mental concerts: musical imagery and auditory cortex. Neuron 47:9–12.

